# Acute Toxicological Profile of Pharmaceutical-Grade Nicotinamide Riboside: A Route-Dependent Assessment Across Intravenous, Intramuscular, and Subcutaneous Administration

**DOI:** 10.64898/2026.02.27.708010

**Authors:** Jun Kwon, Yasmeen Nkrumah-Elie, Jolie Sima Mavoyan, Manjunath DB, Harish AN, Andrew Shao

**Affiliations:** ChromaDex, Inc., a Niagen Bioscience company, Los Angeles, California, United States; Adgyl Lifesciences Private, Ltd., Bengaluru, India

**Keywords:** NAD+, nicotinamide riboside, intravenous, intramuscular, subcutaneous

## Abstract

Nicotinamide riboside chloride (NR-Cl) has been studied predominantly by the oral route, while information regarding its toxicity following parenteral administration is limited. To characterize route-dependent acute toxicity and estimate median lethal doses (LD_50_), pharmaceutical-grade NR-Cl was evaluated following bolus intravenous (IV), intramuscular (IM), and subcutaneous (SC) administration in female Sprague Dawley rats, in three independent studies. All studies were conducted using an adapted OECD Guideline 425 Up-and-Down procedure, modified for parenteral administration, in the absence of standardized route-specific OECD guidance. Animals received a single dose of NR-Cl via the respective administration route and were monitored for mortality, clinical signs, bodyweight changes, and gross pathological findings over a 14-day observation period. Following IM and SC administration, no mortalities were observed at doses up to 2000 mg/kg, and LD_50_ values for both routes were determined to be greater than 2000 mg/kg. In contrast, IV administration yielded an estimated LD_50_ of approximately 2000 mg/kg. These findings demonstrate that the acute toxicity of NR-Cl differs by route of administration and establish foundational safety benchmarks to support future research.

## 1 Introduction

Nicotinamide riboside (NR) is a pyridine-nucleoside classified as member of the vitamin B_3_ family due to its biochemical activity as a precursor to nicotinamide adenine dinucleotide (NAD+), a coenzyme central to cellular energy metabolism. The patented chloride salt form of NR (NR-Cl, available as Niagen^®^) has undergone extensive safety testing for oral use and is Generally Recognized as Safe (GRAS) notified as a food ingredient to the U.S. Food and Drug Administration (FDA) (FDA 2016). The delivery of oral NR-Cl is characterized by intestinal absorption and first-pass hepatic metabolism, yielding systemic exposure to NR and related metabolites such as nicotinic acid and nicotinamide (Yaku et al. 2025). While oral delivery has been shown to elevate circulating and tissue NAD+ concentrations in both humans and rodent models (Trammell et al. 2016; Conze et al. 2019; Christen et al. 2026), parenteral administration may offer clinical utility in certain scenarios due to its ability to bypass gastrointestinal and hepatic first-pass metabolism effects, and thereby potentially greater skeletal muscle bioavailability (Liu et al. 2018; Damgaard et al. 2022). The bioavailability limitations of orally administered NR-Cl require daily administration of 10-14 days to reach steady-state levels of NAD+ (Conze et al. 2019; Yaku et al. 2025)), whereas IV administration assumes immediate 100% bioavailability, presumably resulting in rapid benefits (Hawkins et al. 2024; Reyna et al. 2026).

Initial toxicological assessment of a compound includes the determination of the median lethal dose (LD_50_) to establish acute safety margins and inform dose selection for subsequent subchronic and chronic repeat-dose studies. It also informs the estimation of the maximum tolerable or safe dose through the inclusion of safety factors and calculations of the human equivalent dose (HED). The Organization for Economic Co-operation and Development (OECD) Guideline for the Testing of Chemicals No. 425 provides a validated framework for acute oral toxicity testing using the Up-and-Down Procedure (UDP), which minimizes the number of animals required while maintaining statistical and methodological rigor to estimate the LD_50_ of the test chemical (OECD 2022). Though the OECD 425 procedures are designed specifically for oral gavage and do not address parenteral routes, adaptation of this methodology to intravenous, intramuscular, and subcutaneous routes allows for a systematic characterization of route-dependent toxicity profiles.

While published acute toxicity data for NR-Cl have described oral LD_50_ values exceeding 5000 mg/kg in rodents (Conze et al. 2016), comparative data for parenteral routes remain absent from the literature. Given the potentially unique pharmacokinetic and toxicokinetic profiles associated with intravenous and injection delivery, extrapolation from oral toxicity data may not sufficiently predict parenteral safety. Therefore, the present study aimed to determine the LD_50_ of pharmaceutical-grade NR-Cl, the active pharmaceutical ingredient of NiagenPlus^®^, following three routes of administration – intravenous (IV), intramuscular (IM), and subcutaneous (SC). The study design adhered to an adapted version of the OECD Test Guideline 425 modified for parenteral use, employing female Sprague Dawley rats, and a systematic evaluation of mortality, clinical signs, bodyweight changes, and gross pathology. These findings provide foundational toxicological data regarding the parenteral use of NR-Cl and inform the development of future repeat-dose studies.

## 2 Materials and methods

### 2.1. Test articles

The test article, pharmaceutical-grade β-nicotinamide riboside chloride (pyridinium, 3- (aminocarbonyl)-1-beta-D-ribofuranosyl-chloride; CAS Number 23111-00-4), trade name NiagenPlus^®^, was supplied by ChromaDex, Inc., a Niagen Bioscience company (Los Angeles, CA) and manufactured by Dr. Reddys Laboratories, Ltd. (Telangana, India). The material was provided as a white crystalline powder with certificates of analyses reporting lot purities of 98.9% by weight as determined by high-performance liquid chromatography (Batch Lot ACRH002700, manufactured August 3, 2024; retest date August 2, 2025). Identity and purity were verified through spectroscopic and chromatographic methods. The test article was stored under refrigerated conditions (2-8°C) and formulated freshly on the day of administration by dissolution in 0.9% normal saline.

### 2.2. Median lethal dose (LD_50_) studies

Three independent acute toxicity experiments were performed at Adgyl Lifesciences Private, Ltd., (Bengaluru, India), an AAALAC-accredited facility operating under Good Laboratory Practice (GLP) conditions. Studies were conducted between May and September 2025, and received approval from the Institutional Animal Ethics Committee (Proposal No. 092/Oct-024) in accordance with the Committee for the Control and Supervision of Experiments on Animals (CCSEA) guidelines, Government of India.

All studies were conducted in healthy female Sprague-Dawley rats (*Rattus norvegicus*; 8-12 weeks of age; 210-277 g bodyweight at treatment), obtained from Hylasco Biotechnology Pvt Limted (India). In accordance with general recommendations described in the OECD 425 guidelines, female rats were selected on the basis that they tend to be slightly more sensitive in conventional LD_50_ tests in cases where differences are observed, though there is typically little difference in sensitivity between sexes. At the time of selection, bodyweight variation did not exceed ±20% of the mean for previously dosed animals. All animals were nulliparous and non-pregnant. Rats were housed individually in polysulfone cages with steam sterilized corn cob bedding under controlled standard environmental conditions (temperature: 20-24 ° Celsius; relative humidity: 55-64%; 12-hour light-dark cycle). Animals received standard pellet feed, manufactured by Krishna Valley Agro Tech LLP (Sangli, Maharashtra, India) and purified water, ad libitum. Acclimatization periods lasted from 5 to 21 days. Each rat was identified by cage card and turmeric color body marking, with crystal violet used for temporary body marking during the acclimatization period.

The acute toxicity testing approach was adapted from the OECD Guideline for Testing of Chemicals, No. 425 (Acute Oral Toxicity – Up-and-Down Procedures, adopted October 16, 2008 and corrected June 30, 2022). In the absence of route-specific testing guidance, these guidelines, originally designed for oral administration, were modified for parenteral administration – intravenous, subcutaneous, and intramuscular injections. The Up-and-Down Procedure involves sequential dosing of individual animals with approximate 48-hour intervals between doses, with subsequent dose levels determined based on survival outcomes using AOT425 statistical methods. Dosing continues until stopping criteria are met, defined as three consecutive survivals at the limit dose or the derivation of an LD_50_ estimate. The AOT425StatPgm software was used for the selection of doses and LD_50_ calculations. As specified in the guidelines, animals were observed for 14 days following dose administration, with animals found dead during this period necropsied immediately, and all surviving animals euthanized and necropsied at the conclusion of the observation period. Although OECD 425 guidelines include a limit test for substances anticipated to have low toxicity, a limit test design (i.e., initiation at 2000 mg/kg) was not employed in the present investigation. A general lack of existing data on the toxicology of parenterally administered NR or related derivative compounds in the published literature precluded the anticipation of safety at 2000 mg/kg; therefore, a main test approach was used across all routes to allow characterization of potential dose-dependent acute effects. An initial dose of 650 mg/kg NR-Cl was selected, with a dose progression factor of 1.5 (corresponding to a slope of 7; antilog of 0.14), and a final possible dose of 2000 mg/kg (limit dose). In the event of any mortality at the initial dose, subsequent dosing was to proceed at lower levels accordingly, i.e., the initial dose divided by 1.5.

All dose formulations were prepared prior to the administration by dissolving weighed quantities of NR-Cl in normal saline 0.9% w/v until the desired test item concentrations were achieved. Formulations were prepared aseptically under a biosafety cabinet using sterile glassware. Beakers were pre-calibrated by marking the meniscus level corresponding to the final volume. Stability and homogeneity of NR-Cl in 0.9% normal saline were previously assessed for stability and homogenous solution formation, thus repeat characterization was not performed in the present studies. This approach represents a documented exception to the OECD principles of GLP; this was justified on the basis that formulations were prepared using methods identical to those employed in the prior stability assessments, and that the test material was adequately mixed and administered immediately following the formulation.

Following dosing, animals were observed for clinical signs on Day 1 (day of administration) at repeated intervals, followed by once-daily observations through Day 15. Observations included assessment of skin, fur, eyes, mucous membranes, respiratory and circulatory systems, central and autonomic nervous system function, somatomotor activity, and behavior. Bodyweights were recorded on Day 1, Day 8, and Day 15. Animals surviving to the end of the observation period were euthanized by isoflurane anesthesia and subjected to detailed necropsy. Gross pathological findings were recorded for all animals.

#### 2.2.1. Intramuscular (IM)

In this study, each animal received a single intramuscular dose of 0.5 mL/kg solution, administered in equal volumes bilaterally into the left and right quadriceps femoris muscles using a 26-gauge, ½-inch needle. Hair at the injection site was clipped and the injection sites were marked for each animal. Dosing was initiated at 650mg/kg bodyweight, while subsequent animals received doses of 910, 1260, 2000mg/kg bodyweight. Dosing was stopped once three consecutive rats survived at the 2000mg/kg dose, in accordance with the stopping criteria.

#### 2.2.2. Intravenous (IV)

Intravenous administration was performed via a single tail vein injection using a 27-gauage,1/2-inch needle attached to a disposable plastic syringe. The injection volume was 5 ml/kg bodyweight, administered as a single bolus injection. Dosing was initiated at 650 mg/kg, subsequently followed by 910, 1260, and 2000 mg/kg based on sequential survival outcomes. Dosing continued according to the Up-and-Down procedure until stopping criteria were met following repeated survival at 1260 mg/kg after mortality was observed at 2000 mg/kg.

#### 2.2.3. Subcutaneous (SC)

Subcutaneous administration was performed via injection into the subcutaneous space of the cervical region. The total volume of 5 ml/kg was equally divided between two sites (2.5 ml/kg/site) using a 27-gauge, ½-inch needle attached to a disposable plastic syringe. Hair was clipped at injection site prior to administration, and sites were marked. The dose-escalation protocol mirrored that of the intramuscular study: starting at 650 mg/kg, with escalation to 910, 1260, and 2000 mg/kg, with termination after three consecutive survivals at the limit dose.

### 2.3. Statistical Analysis

The LD_50_ estimate and 95% confidence interval were calculated using AOT425StatPgm software based on the sequence of doses and mortality outcomes.

## 3 Results

### 3.1 Intramuscular

Divided single dose IM administration of NR-Cl produced no mortality and clinical signs at any of the tested dose levels. Necropsy examination revealed no treatment-related lesions or abnormalities. The summary of outcomes is presented in Table 1.

**Table 1.**
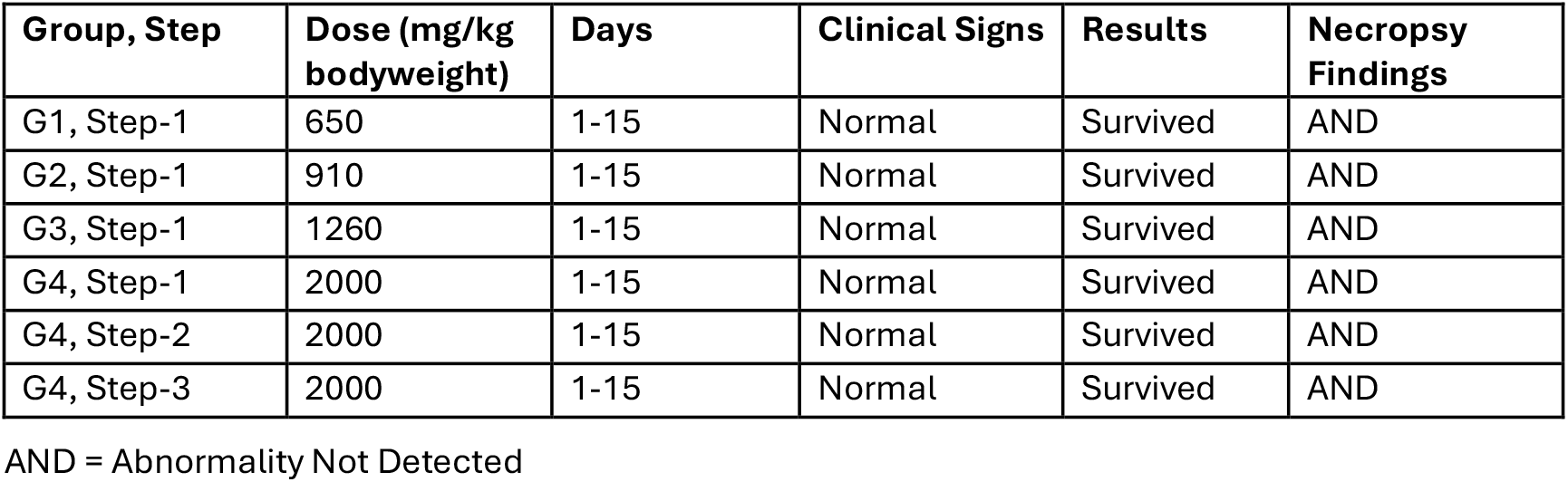
Intramuscular dose administration, clinical signs and necropsy findings in female Sprague-Dawley Rats.

| Group, Step | Dose (mg/kg bodyweight) | Days | Clinical Signs | Results | Necropsy Findings |
| --- | --- | --- | --- | --- | --- |
| G1, Step-1 | 650 | 1-15 | Normal | Survived | AND |
| G2, Step-1 | 910 | 1-15 | Normal | Survived | AND |
| G3, Step-1 | 1260 | 1-15 | Normal | Survived | AND |
| G4, Step-1 | 2000 | 1-15 | Normal | Survived | AND |
| G4, Step-2 | 2000 | 1-15 | Normal | Survived | AND |
| G4, Step-3 | 2000 | 1-15 | Normal | Survived | AND |

AND = Abnormality Not Detected

Bodyweight changes were monitored throughout the treatment period. Treatment-related effects on bodyweight were not observed at any of the tested dose levels. The rats gained bodyweight over the course of the observation period.

On the basis of these findings, the IM LD_50_ was determined to exceed 2000 mg/kg. The No Observed Adverse Effect Level (NOAEL) was ≥2,000 mg/kg, as no adverse effects were observed at any tested dose including the limit dose.

### 3.2. Intravenous

IV administration of NR-Cl resulted in mortality at 2000 mg/kg. The summary of outcomes is presented in Table 2.

**Table 2.** Intravenous dose administration, clinical signs and necropsy findings in female Sprague-Dawley Rats.

| Group, Step | Dose (mg/kg bodyweight) | Day(s) | Clinical Signs | Results | Necropsy Findings |
| --- | --- | --- | --- | --- | --- |
| G1, Step-1 | 650 | 1-15 | Normal | Survived | AND |
| G2, Step-1 | 910 | 1-15 | Normal | Survived | AND |
| G3, Step-1 | 1260 | 1 | Tail discoloration (brown color) | Survived | AND |
|  |  | 2-15 | Tail discoloration (black color) |  |  |
|  |  | 12-15 | Tail detachment |  |  |
| G4, Step-1 | 2000 | 1 | Tail discoloration (brown color) | Survived | AND |
|  |  | 1-15 | Tail discoloration (black color) |  |  |
|  |  | 9-15 | Tail detachment |  |  |
| G4, Step-2 | 2000 | 1 | Posture recumbent | Died | AND |
| G3, Step-2 | 1260 | 1 | Tail discoloration (brown color) | Survived | AND |
|  |  | 2-15 | Tail discoloration (black color) |  |  |
| G4, Step-3 | 2000 | 1 | Posture recumbent | Died | AND |
| G3, Step-3 | 1260 | 1 | Tail discoloration (brown color) | Survived | AND |
|  |  | 2-15 | Tail discoloration (black color) |  |  |
|  |  | 12-15 | Tail detachment |  |  |

AND = Abnormality Not Detected

At 650 and 910 mg/kg (G1 Step 1; G2 Step 1), no clinical signs or mortalities were observed. The animals survived through Day 15 with no evidence of bodyweight suppression. At 1260 mg/kg (G3 Step 1), clinical signs of tail discoloration were noted, with the normal pink/flesh coloration changing to brown (Day 1) and progressing to black (Days 2-15), ultimately followed by tail detachment from Days 12-15 - the animal survived through Day 15.

At 2000 mg/kg (G4 Step 1), both tail discoloration (brown on day 1, black Days 1-15) and tail detachment (Days 9-15) were observed; the animal survived. At the subsequent 2000 mg/kg dose (G4 Step 2), posture recumbency occurred within 1 minute of administration, followed by immediate death, prompting a dose reduction to 1260 mg/kg (G3 Step 2), at which the animal survived. The dose was then increased to 2000 mg/kg (G4 Step 3), where posture recumbency was again observed within 1 minute, followed by immediate death. The dose was subsequently reduced to 1260 mg/kg (G3 Step 3), where the animal survived. Similar tail discoloration and detachment patterns observed earlier were once again noted during G3 Steps 2 and 3, both of which survived. Dosing was terminated after three consecutive survivals at 1260 mg/kg, consistent with the stopping criteria defined in the OECD guidelines. A total of 8 animals were used across the sequential dosing steps.

Bodyweights at baseline ranged from 250.84 to 276.78 g and progressed without evidence of treatment-related effects at all dose levels, with the exception of a slight reduction observed at 2000 mg/kg (Supplemental Taable 2).

Necropsy of animals that died at 2000 mg/kg revealed no abnormalities detected. Surviving animals showed tail discoloration and tail detachment at 1260 and 2000 mg/kg, consistent with in-life observations. No systemic gross pathological findings were observed.

The calculated IV LD_50_ was 2000 mg/kg (95% confidence interval: 1319-5910 mg/kg) based on AOT425 software analysis. The sponsor proposed a more conservative LD_50_ estimate of 1260 to 2000 mg/kg. The physiological cause of death was not determined, as necropsy results indicated no abnormalities. A NOAEL of 910 mg/kg was selected, representing the highest dose at which no mortality, clinical signs, or pathological findings were observed.

### 3.3 Subcutaneous

SC administration of NR-Cl produced no mortality at any tested dose. The summary of outcomes is presented in Table 3.

**Table 3.**
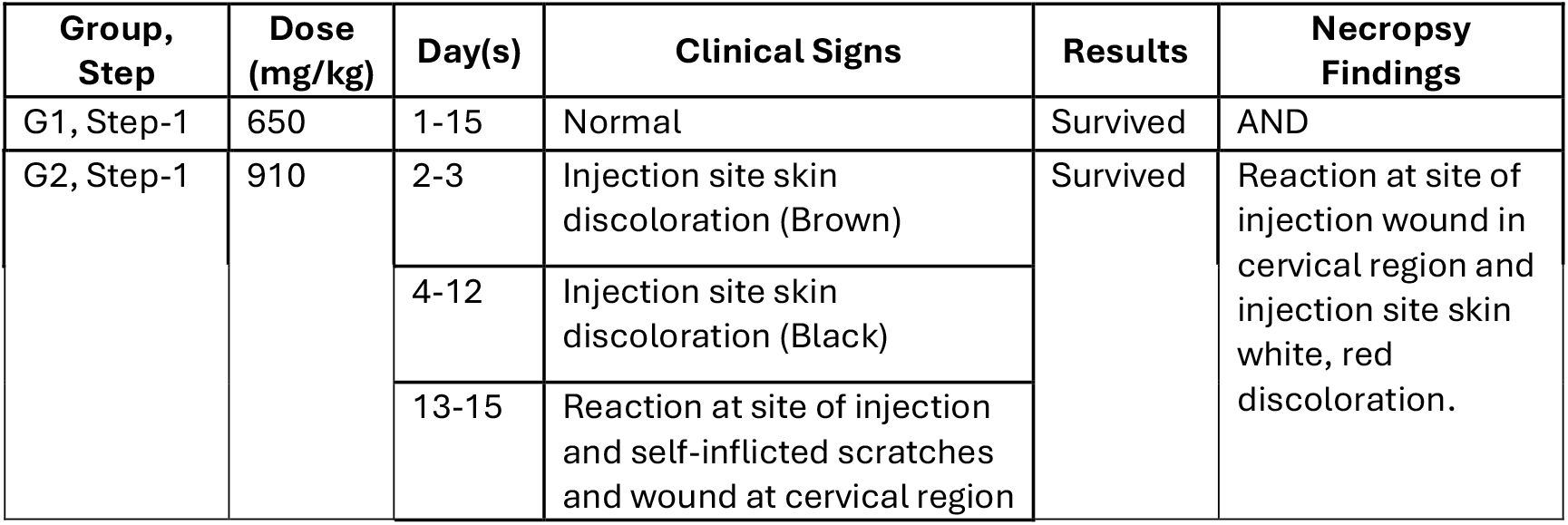
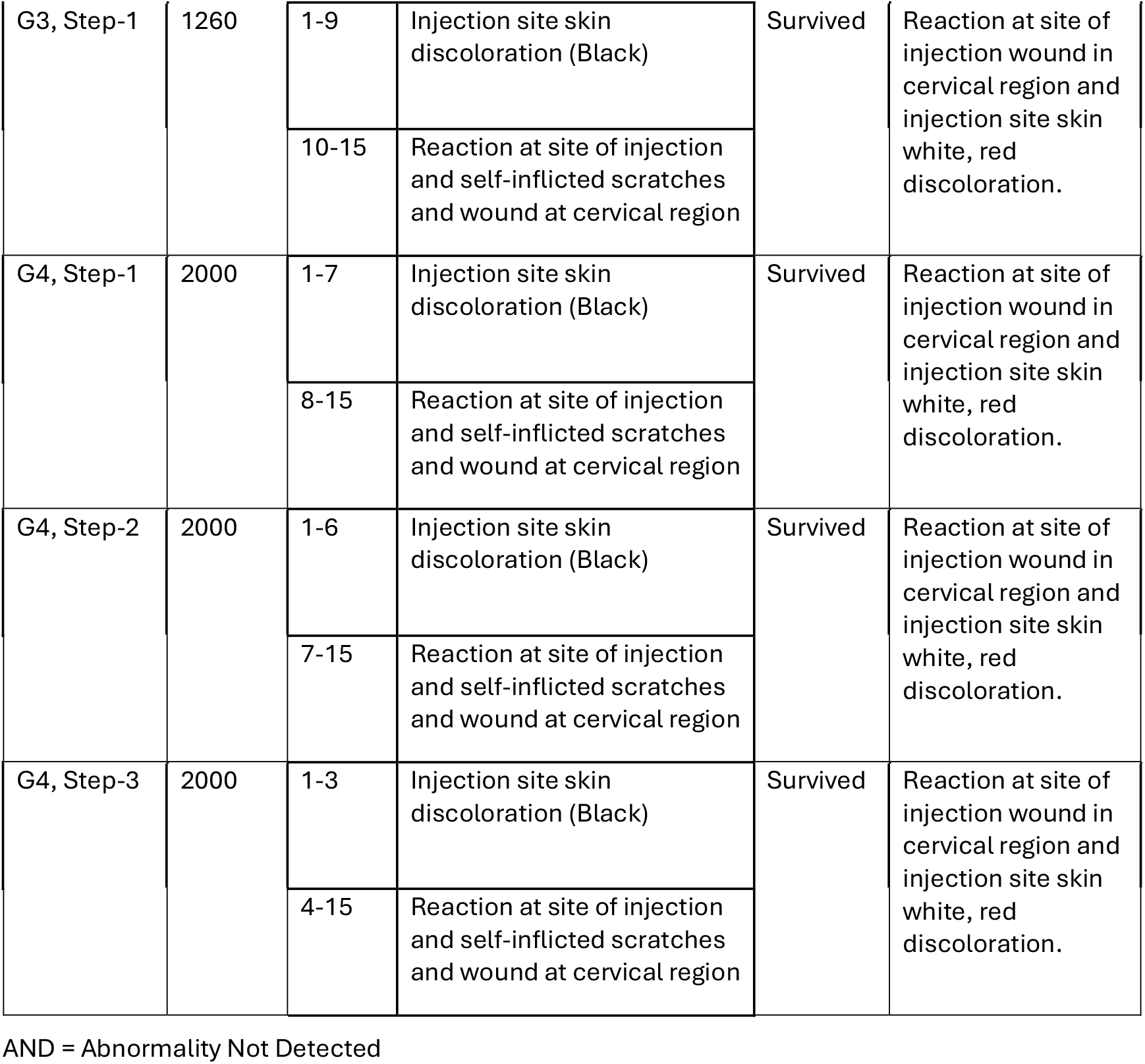
Subcutaneous dose administration, clinical signs and necropsy findings in female Sprague-Dawley Rats.

| Group, Step | Dose (mg/kg) | Day(s) | Clinical Signs | Results | Necropsy Findings |
| --- | --- | --- | --- | --- | --- |
| G1, Step-1 | 650 | 1-15 | Normal | Survived | AND |
| G2, Step-1 | 910 | 2-3 | Injection site skin discoloration (Brown) | Survived | Reaction at site of injection wound in cervical region and injection site skin white, red discoloration. |
|  |  | 4-12 | Injection site skin discoloration (Black) |  |  |
|  |  | 13-15 | Reaction at site of injection and self-inflicted scratches and wound at cervical region |  |  |
| G3, Step-1 | 1260 | 1-9 | Injection site skin discoloration (Black) | Survived | Reaction at site of injection wound in cervical region and injection site skin white, red discoloration. |
|  |  | 10-15 | Reaction at site of injection and self-inflicted scratches and wound at cervical region |  |  |
| G4, Step-1 | 2000 | 1-7 | Injection site skin discoloration (Black) | Survived | Reaction at site of injection wound in cervical region and injection site skin white, red discoloration. |
|  |  | 8-15 | Reaction at site of injection and self-inflicted scratches and wound at cervical region |  |  |
| G4, Step-2 | 2000 | 1-6 | Injection site skin discoloration (Black) | Survived | Reaction at site of injection wound in cervical region and injection site skin white, red discoloration. |
|  |  | 7-15 | Reaction at site of injection and self-inflicted scratches and wound at cervical region |  |  |
| G4, Step-3 | 2000 | 1-3 | Injection site skin discoloration (Black) | Survived | Reaction at site of injection wound in cervical region and injection site skin white, red discoloration. |
|  |  | 4-15 | Reaction at site of injection and self-inflicted scratches and wound at cervical region |  |  |

AND = Abnormality Not Detected

At 650 mg/kg (G1 Step 1), no clinical abnormalities were detected throughout the observation period. At 910 mg/kg (G2 Step 1), injection site skin discoloration (brown, Days 2-3; black, Days 4-12) was observed, with reaction at site of injection wound and self-inflicted scratches in the cervical region from Days 13-15.

At 1260 mg/kg (G3 Step 1), injection site skin discoloration (black, Days 1-9) was observed, with reaction at site of injection wound and self-inflicted scratches in the cervical region from Days 10-15.

At 2000 mg/kg (G4 Step 1), injection site skin discoloration (black, Days 1-7) was noted, with reaction at the site of injection wound and self-inflicted scratches in the cervical region from Days 8-15. At 2000 mg/kg (G4 Step 2), injection site skin discoloration (black, Days 1-6) occurred with wound and scratches from Days 7-15. At 2000 mg/kg (G4 Step 3) injection site skin discoloration (black, Days 1-3) was observed, with reaction at site of injection wound and self-inflicted scratches from Days 4-15.

Bodyweights at baseline ranged from 237.51 to 248.79 g. Bodyweight progression remained stable across all dose groups, with no treatment-related effects observed.

Gross necropsy findings were limited to injection site alterations. At 650 mg/kg, no abnormalities were detected. At 910, 1260, and 2000 mg/kg, necropsy revealed reaction at site of injection wound in the cervical region and injection site skin white and red discoloration. No systemic pathology was observed.

On the basis of these findings, the SC LD_50_ was determined to exceed 2000 mg/kg. A NOAEL of 650 mg/kg was specified, the dose at which no mortality or injection site reactions were observed.

### 3.4 Human Equivalent Dose Translation

Translation of rat LD_50_ values to human equivalent doses (HED) was performed using the body surface area scaling method adopted by the FDA, which provides a preliminary basis for clinical risk assessment (FDA 2005). The conversion employed species-specific Km factors (rat = 6, adult human = 37), and used the following formula: HED (mg/kg) = Animal dose (mg/kg) x (Animal Km / Human Km). Absolute doses for a 70-kg adult human were calculated by multiplying the derived HED values by 70 kg. Results are presented in Table 4.

**Table 4.** Human equivalent doses from female Sprague-Dawley Rats.

| Route of administration | Dose (mg/kg) | Observation | HED (mg/kg) | HED for a 70-kg individual (g) |
| --- | --- | --- | --- | --- |
| Intramuscular | >2000 | LD<sub>50</sub> (limit not reached) | >324 | 23 |
|  | 650-2000 | No mortality, no adverse findings (NOAEL at the highest dose) | 105-324 | 7-23 |
| Subcutaneous | >2000 | LD<sub>50</sub> (limit not reached) | >324 | 23 |
|  | 650 | Injection site reactions absent (NOAEL) | 105 | 7 |
|  | 910-2000 | Injection site reactions present, no mortality | 147-324 | 10-23 |
| Intravenous | 2000 | LD<sub>50</sub> (calculated), 2/3 animals died | 324 | 23 |
|  | 1260-2000 | Conservative LD<sub>50</sub> range | 204-324 | 14-23 |
|  | 1260 | Lower doses at which tail lesions were observed, no mortality | 204 | 14 |
|  | 650-910 | No mortality, no tail lesions (NOAEL = 910 mg/kg) | 105-147 | 7-10 |

## 4 Discussion

The present three-part investigation provides systematic acute toxicity data for pharmaceutical-grade NR-Cl administered via three parenteral routes in female Sprague-Dawley rats using a sequential Up-and-Down design. The findings demonstrate route-dependent differences in lethality and local tolerability. IV administration yielded an estimated LD_50_ of 2000 mg/kg (conservative range: 1260-2000 mg/kg) with mortality occurring in 2 of 3 animals at the limit dose. On the other hand, IM and SC routes were well-tolerated at all tested doses up to 2000 mg/kg, with LD_50_ values exceeding the limit dose. These results indicate that both dose and route of administration modulate acute NR-Cl toxicity.

The occurrence of mortality at 2000 mg/kg following IV administration, in contrast to the absence of mortality at the same dose following IM and SC administration, may at least in part be explained by its direct delivery to systemic circulation, likely resulting in higher peak concentrations, and 100% bioavailability. The rapid onset of recumbency, in the absence of treatment-related gross-pathological lesions, followed by immediate death observed at 2000 mg/kg may be suggestive of an acute systemic effect associated with accelerated peak exposure, though this cannot be confirmed without pharmacokinetic data. The mortalities may also in part reflect hemodynamic consequences associated with rapid, high-concentration bolus delivery. The dose formulations of 1260 and 2000 mg/kg required NR-Cl concentrations of 252 and 400 mg/ml, respectively, which may have exceeded the osmotic tolerance of the vasculature. The absence of tail lesions of animals receiving lower doses of 650 and 910 mg/kg, corresponding to volumes of130 and 182 mg/mL, delivered at equivalent volumes of 5 ml/kg, lends further support for a contributary role of concentration, rather than volume, on the observed effects.

By comparison, IM and SC administration were not associated with systemic clinical signs and no mortality at doses up to 2000 mg/kg. These routes permit absorption-limited, gradual systemic exposure, lower peak concentrations, and longer time to maximum exposure compared to IV bolus delivery, despite equivalent administered doses. Both IM and SC doses were divided between two anatomical sites on the animal, administered seconds apart. The complete absence of mortality at 2000 mg/kg with SC and IM routes, compared to 2/3 mortality at the same dose given IV, demonstrates that absorption kinetics are one of the determinants of acute NR-Cl lethality.

A similar pattern emerges in relation to local findings. Following IV dosing, the consistent observation of tail discoloration and subsequent detachment at 1260 and 2000 mg/kg in surviving animals. These effects were limited to the injection site and likely reflect localized vascular injury associated with tail vein administration. In the SC study, injection site reactions included discoloration, wounds, and self-inflicted lesions at doses of 910 mg/kg and above, persisting throughout the observation period. In the absence of systemic pathological findings, these findings also likely indicate localized tissue irritation. Conversely, no adverse consequences, whether systemic or local, were observed following IM administration under the conditions tested, suggesting a substantial margin of safety. The smaller injection volume employed for IM dosing (0.5 ml/kg total) relative to SC (5 ml/kg total), however, limits direct head-to-head comparisons between these two routes.

The human equivalent dose (HED) translations provide a framework evaluating the potential clinical relevance of these findings (FDA 2005). For IM and SC routes, where rat LD_50_ values exceeded 2000 mg/kg, the HED would exceed 324 mg/kg, corresponding to >23 g for a 70-kg individual. Concerning the SC route, injection site reactions were absent at 650 mg/kg (equivalent to 105 mg/kg, or 7 g for a 70-kg individual), but present at 910 mg/kg and above (147 mg/kg or 10 g and higher for a 70-kg individual) – suggesting a threshold dose for local tissue irritation. Of note, however, is that in the present study, IM and SC doses were divided between two injection sites to minimize local tissue burden and accommodate volume constraints. On the other hand, in clinical practice, injections are typically delivered as a single bolus dose at a single site. This methodological difference implies that the threshold for injection site reactions in humans may be lower than the HED calculations imply, as concentrating the full dose at a single anatomical location would increase local tissue exposure compared to the distributed dosing method used in this study.

Concerning IV administration, the present study reported a conservative rat LD_50_ range of 1260-2000 mg/kg, which translates to human equivalent doses of 204-324 mg/kg, or 14-23 g for a 70-kg individual. The lower doses at which tail lesions were observed intravenously (1260 mg/kg, equivalent to 204 mg/kg or 14 g in humans) and the doses without mortality (650-1260 mg/kg in rats, equivalent to 105 -204 mg/kg, or 7-14 g in humans) provide additional reference points for evaluating vascular tolerability. Of note, however, is that the experimental design employed a single bolus injection via the tail vein, delivering the entire dose within seconds. This is in contrast to clinical practice, which generally involves infusion over minutes to hours rather than bolus administration. Indeed, early clinical work by Hawkins et al., who used an adaptive approach wherein the participants self-selected a tolerable infusion rate, documented mean NR-Cl IV infusion times of roughly 25 minutes for a 500 mg dose (Hawkins et al., 2024). Therefore, in a real-world clinical setting, infusion-based delivery would likely result in an attenuation of peak plasma concentrations and reductions in vascular stress, compared to the rapid bolus delivery employed in the present study. Future studies are needed to systematically confirm whether a slower rate of IV delivery, as is often employed in clinical practice to preserve tolerability, mitigates the onset of injection site damage and mortality.

The No Observed Adverse Effect Levels (NOAELs) identify dose thresholds at which no adverse findings are observed. The NOAEL specified for the IM route was the highest, at ≥2000 mg/kg (HED ≥324 mg/kg or 23 g), as no adverse effects or moralities were observed at any of the tested doses. SC administration similarly did not produce mortalities at any of the dose levels, however, the NOAEL corresponded to the lowest dose tested of 650 mg/kg (HED 105 mg/kg or 7 g), at which injection site reactions were absent. The IV administration resulted in mortality at the limit dose, and a NOAEL of 910 mg/kg (HED: 147 mg/kg or 10 g for a 70-kg individual), as no clinical signs were observed at this level of exposure.

Collectively, these findings hold the potential to inform formulation development and clinical dosing strategies. The HED calculations offer a useful starting point for approximating equivalent human exposure, representing initial estimates derived from the normalization of doses to body surface area. However, they are not intended to serve as definitive clinical dose predictions. Interspecies differences in NAD+ metabolism, vascular physiology, dermal tolerance and distribution kinetics introduce uncertainty, as do methodological differences between the experimental bolus/divided-site administration employed here and the slower infusion or single-site injection practices typical in clinical practice. Therefore, the presented HED values should be interpreted as preliminary benchmarks.

The present study also introduces questions that warrant further investigation through pharmacokinetic and safety studies. Studies incorporating dose escalation and comparing bolus versus slow infusion of NR-Cl IV at equivalent total doses, would clarify the effects of dose and time course of delivery on tolerability. Moreover, pharmacokinetic studies of parenteral doses are needed to determine if mortality or the onset of adverse effects correlate with peak plasma concentrations of NR, or its metabolites including NAD+, nicotinamide, and related derivatives. Patient-specific characteristics and preferences may also inform the appropriate route of parenteral NR-Cl administration. Route-comparison studies across multiple therapeutic drugs have demonstrated that factors such as BMI, age, and medical comorbidities may modify the relative efficacy and safety of different parenteral routes, in certain cases. For instance, the administration of hydrocortisone via the SC route demonstrates BMI-dependent pharmacokinetics, whereas no such relationship is observed for the IM route (Jin et al. 2015). Whether similar variations in uptake or responses apply to NR-Cl is unknown at the present time; therefore, this represents a relevant topic of future study to eventually inform dose selection or route optimization in clinical practice.

Several methodological considerations warrant discussion. The adaptation of OECD 425 guidelines from oral to parenteral administration is pragmatic and preserves the statistical framework of the Up-and-Down procedures. Nonetheless, it is worth acknowledging that the underlying assumptions built into the test are based on absorption kinetics developed for oral routes. The 48-hour intervals between doses and observation periods are appropriate for detecting acute effects. The use of female rats aligns with guideline recommendations, in addition to prior literature indicating greater sensitivity of females to NR-Cl, though these studies were addressing oral administration. Finally, though the stability and homogeneity of NR-Cl in formulation were not reassessed in the present studies, these parameters have been previously characterized elsewhere, documenting stable and homogenous solution formation under identical preparation conditions.

## 5 Conclusion

In conclusion, the acute toxicity of pharmaceutical-grade NR-Cl varies by mode of administration in female Sprague-Dawley rats. IV administration yielded an LD_50_ of approximately 2000 mg/kg, with rapid onset mortality and dose-dependent tail lesions, however, without evidence of systemic toxicity. SC and IM routes resulted in LD_50_ in excess of 2,000 mg/kg. IM administration was well-tolerated at all dose levels up to 2000 mg/kg, producing no mortalities or adverse clinical findings. SC administration produced no mortality at doses up to 2000 mg/kg but elicited local injections site reactions starting at 910 mg/kg, without systemic consequences. Taken together, these data suggest a concentration-driven adverse effect on the local site of injection, rather than an inherent toxic effect of NR-Cl itself. In addition, the adverse vascular and mortality effects of IV administration may also, in part, be explained by the rapid bolus delivery. The present study provides foundational toxicological information for the evaluation of parenteral NR-Cl. Moreover, the differential toxicological profiles across intravenous, intramuscular, and subcutaneous routes underscore the importance and necessity of route-specific safety assessments.

## 7 Author Contributions

Conceptualization and methodology, JK, YNE, AS; writing, JK, JSM; YNE; supervision: YNE. All authors contributed to the development and revision of the manuscript.

All experiments, data curation, and statistical analyses were performed at Adgyl Lifesciences Private, Ltd.

## 9 Acknowledgments

The authors would like to thank the staff at Adgyl Lifesciences Private, Ltd for conducting the laboratory experimental work.

## 11 Declaration of Generative AI Use

Generative AI Use was not used in the development of this manuscript

