## Supplementary Tables for "Acute Toxicological Profile of Pharmaceutical-Grade Nicotinamide Riboside: A Route-Dependent Assessment Across Intravenous, Intramuscular, and Subcutaneous Administration"

Supplementary table 1. Intramuscular dose administration, body weight observations in female Sprague-Dawley Rats

| Group, Step | Dose (mg/kg body weight) | Body Weight (g) | | | | |
| --- | --- | --- | --- | --- | --- | --- |
|  |  | Initial (Day 1) | Day 8 | Weight change (Day 8 – Initial) | Day 15 | Weight change (Day 15 – Initial) |
| G1, Step-1 | 650 | 230.14 | 256.19 | 26.05 | 287.34 | 57.20 |
| G2, Step-1 | 910 | 233.68 | 260.90 | 27.22 | 279.45 | 45.77 |
| G3, Step -1 | 1260 | 224.46 | 249.48 | 25.02 | 280.39 | 55.93 |
| G4, Step-1 | 2000 | 222.56 | 240.39 | 17.83 | 267.70 | 45.14 |
| G4, Step-2 | 2000 | 220.49 | 246.87 | 26.38 | 287.55 | 67.06 |
| G4, Step-3 | 2000 | 210.56 | 239.20 | 28.64 | 277.31 | 66.75 |

Supplementary table 2. Intravenous dose administration, body weight observations in female Sprague-Dawley Rats

| Group, Step | Dose (mg/kg body weight) | Body Weight (g) | | | | | | |
| --- | --- | --- | --- | --- | --- | --- | --- | --- |
|  |  | Initial (Day 1) | Day 8 | Weight change (Day 8 – Initial) | Day 15 | At death | Weight change (Day 15 – Initial) | Weight change (At death – Initial) |
| G1, Step-1 | 650 | 250.84 | 271.12 | 20.28 | 284.02 | NA | 33.18 | NA |
| G2, Step-1 | 910 | 254.72 | 256.84 | 2.12 | 288.56 | NA | 33.84 | NA |
| G3, Step-1 | 1260 | 260.95 | 267.99 | 7.04 | 269.56 | NA | 8.61 | NA |
| G4, Step-1 | 2000 | 275.62 | 262.23 | -13.39 | 276.53 | NA | 0.91 | NA |
| G4, Step-2 | 2000 | 270.35 | NA | NA | NA | 270.09 | NA | -0.26 |
| G3, Step-2 | 1260 | 275.43 | 281.17 | 5.74 | 283.33 | NA | 7.9 | NA |
| G4, Step 3 | 2000 | 276.78 | NA | NA | NA | 276.53 | NA | -0.25 |
| G3, Step 3 | 1260 | 273.82 | 276.78 | 2.96 | 282.54 | NA | 8.72 | NA |

NA = Not applicable

Supplementary table 3. Subcutaneous dose administration, body weight observations in female Sprague-Dawley Rats

| Group, Step | Dose (mg/kg body weight) | Body Weight (g) | | | | |
| --- | --- | --- | --- | --- | --- | --- |
|  |  | Initial (Day 1) | Day 8 | Weight change (Day 8 – Initial) | Day 15 | Weight change (Day 15 – Initial) |
| G1, Step-1 | 650 | 237.51 | 259.13 | 21.62 | 274.69 | 37.18 |
| G2, Step-1 | 910 | 245.34 | 263.85 | 18.51 | 291.67 | 46.33 |
| G3, Step -1 | 1260 | 245.31 | 259.41 | 14.10 | 288.74 | 43.43 |
| G4, Step-1 | 2000 | 240.28 | 265.74 | 25.46 | 290.53 | 50.25 |
| G4, Step-2 | 2000 | 240.73 | 261.56 | 20.83 | 299.76 | 59.03 |
| G4, Step-3 | 2000 | 248.79 | 274.69 | 25.90 | 309.60 | 60.81 |
